# An environmental DNA tool for monitoring the status of the Critically Endangered Smalltooth Sawfish, *Pristis pectinata*, in the Western Atlantic

**DOI:** 10.1101/765321

**Authors:** Ryan N. Lehman, Gregg R. Poulakis, Rachel M. Scharer, Katherine E. Schweiss, Jill M. Hendon, Nicole M. Phillips

## Abstract

The Critically Endangered Smalltooth Sawfish, *Pristis pectinata*, was once widespread in the tropical and subtropical waters of the Atlantic Ocean, but following substantial declines over the past century, the core population is currently confined to southwest Florida in the U.S. and the Bahamas. Recent research and verified public encounter reports suggests that this core population may be stabilizing and, potentially, expanding into formerly occupied areas of their historic range in the Western Atlantic; however, the status of this species in non-core waters is not well understood. Environmental DNA (eDNA) methods provide a relatively cost effective and rapid assessment tool for monitoring species occurrence in aquatic habitats. Here, we have developed an eDNA tool: a species-specific Droplet Digital™ PCR (ddPCR™) assay targeting a 100-base pair portion of the mitochondrial NADH dehydrogenase subunit 2 gene in *P*. *pectinata*, with the ability to reliably detect as little as 0.25 pg of target DNA. The assay was validated by collecting and analyzing a water sample from known *P*. *pectinata* nursery habitat in Florida, which was found to contain an average of 11.54 copies of target DNA/µL (SE = 0.72) in the reaction. The assay was then further tested by placing a juvenile sawfish in an *ex situ* tank and analyzing water samples collected at time intervals. The implementation of this eDNA tool into field surveys will provide additional, reliable data to assess species recovery and aid in prioritizing localities beyond the core range in which to focus research and education initiatives.

## Introduction

Sawfishes are among the most threatened families of marine fishes worldwide (Dulvy et al. 2014), with all five species listed as Critically Endangered or Endangered on the International Union for Conservation of Nature (IUCN) Red List of Threatened Species (Dulvy et al. 2016). All sawfishes have undergone global declines in range and abundance due to direct exploitation, bycatch in fisheries, and habitat loss (Dulvy et al. 2016). These threats are exacerbated by their life history traits (e.g., late maturity, low fecundity, and long life spans), which leave sawfishes susceptible to overexploitation, and makes population recovery a slow process (Stevens et al. 2000; Carlson and Simpfendorfer 2015).

The Critically Endangered Smalltooth Sawfish, *Pristis pectinata*, is thought to have experienced the largest global range contraction of all sawfishes and is currently found in less than 20% of its former range (Dulvy et al. 2016). Once widespread in the tropical and subtropical waters of the Atlantic Ocean, remaining core population(s) are thought to be limited to the U.S. and the Bahamas (Carlson et al. 2013), making these populations of global conservation significance (Dulvy et al. 2016). Within U.S. waters, *P*. *pectinata* were historically found from Texas to the Carolinas (Brame et al. 2019) but saw substantial losses in both range and abundance over the past century, with the current population restricted to southwest Florida (SWFL) by the 1980’s (Norton et al. 2012).

Due to the dramatic declines in range and abundance, *P. pectinata* was listed as Endangered in 2003 under the U.S. Endangered Species Act of 1973 by the National Marine Fisheries Service (NMFS) (NMFS 2003), and a Species Recovery Plan (SRP) was developed to promote recovery and long-term viability of the species in U.S. waters (NMFS 2009; 2018). One characteristic of full species recovery is re-establishment in some or all of the former range (Akçakaya et al. 2018); therefore, the SRP for *P. pectinata* designated 15 recovery regions throughout their historic range in U.S. waters, wherein recovery efforts should occur if species presence is confirmed (NMFS 2009; 2018). As a result of over 15 years of U.S. federal and state protections, scientific advances in the understanding of the biology and ecology of the species, and public education initiatives, the core population of *P. pectinata* in SWFL is believed to be stabilizing (NMFS 2018). One line of evidence for this potential stabilization is the emergence of relatively recent sawfish encounter reports within formerly occupied parts of their historic range in U.S. waters, including in designated recovery regions (NMFS 2018); however, the status of *P. pectinata* in these non-core areas is unknown.

Traditional survey methods for monitoring the status of rare species can be expensive and time-consuming (Lewison et al. 2004). Environmental DNA (eDNA) methods provide a relatively cost effective and rapid assessment tool for monitoring species occurrence in aquatic habitats (Rees et al. 2014; Evans et al. 2017). Water provides a medium for traces of DNA recently shed by organisms (e.g., cellular debris, skin cells, blood, feces, urine), which can be collected and analyzed via genetic assays (Jerde et al. 2011). EDNA has been shown to be as sensitive, and sometimes more sensitive in rare species detections compared to survey methods such as electrofishing (Evans et al. 2017), Baited Remote Underwater Video systems (BRUVs) and underwater visual censuses (UVCs) (Boussarie et al. 2018), and traditional net surveys (Thomsen et al. 2012). EDNA methods also negate the need to capture and handle the target species, making it an ideal tool to assess the presence or absence of a threatened species (Rees et al. 2014). EDNA has been used in targeted, single species detections for a growing number of threatened elasmobranchs, including the Endangered Maugean Skate, *Zearaja maugeana* (Weltz et al. 2017), the Vulnerable Great White Shark, *Carcharodon carcharias* (Lafferty et al. 2018), the Vulnerable Chilean Devil Ray, *Mobula tarapacana* (Gargan et al. 2017), and the Critically Endangered Largetooth Sawfish, *Pristis pristis* (Simpfendorfer et al. 2016).

Here, we develop and validate an eDNA assay to detect the presence of *P*. *pectinata* DNA in water samples, for use as a tool for monitoring their recovery in the Western Atlantic. This tool will allow scientists and managers to better understand the status of *P. pectinata* in non-core areas and provide quantitative baseline data from which to measure progress towards recovery. Such data can also aid in prioritizing recovery regions in which to focus research and education initiatives, playing an important role in adaptive management strategies as *P. pectinata* expands into its former range.

### *Pristis pectinata* eDNA assay development

#### Field and laboratory controls

To reduce the risk of contamination by exogenous DNA or cross-contamination between samples, rigorous controls were used throughout all stages of this research (see Ficetola et al. 2016; Goldberg et al. 2016; Port et al. 2016; Schweiss et al. In press). All water collection bottles and filtering equipment were cleaned prior to each use using a combination of two methods of sterilization; cleaning with 10% bleach followed by either autoclaving at 120°C for 20 min or exposure to UV light for 20 min, depending on the materials. To prevent contamination between the stages of sample processing, water filtration, DNA extractions, and PCR amplifications were conducted in physically isolated laboratories. Furthermore, water samples were filtered in laboratories where contemporary *P. pectinata* tissue had never been present (see Deiner et al. 2015). During water filtration and DNA extraction, designated sterile forceps for each sample were used to handle used filters and gloves were changed between samples to prevent cross-contamination between samples (see Pilliod et al. 2013; Goldberg et al. 2016). During DNA extractions and PCR, aerosol barrier pipette tips were used to prevent cross-contamination between samples (Schweiss et al. In Press). Additionally, no positive samples were included in any PCRs due to the risk of contamination from the positive itself, as per ancient DNA (aDNA) PCR protocols (see Mulligan 2005).

To test for the possibility of contamination, negative control samples were incorporated into water sample collection and each stage of laboratory processing and analyzed through to PCRs, which were conducted in replicates of five (Jerde et al. 2011; Bakker et al. 2017). To test for contamination in the field, 3 L of autoclaved DI water were brought onto the boat and stored in three sterile 1 L Nalgene^®^ bottles on ice until filtration. To test for contamination during filtration, 3 L of autoclaved DI water were filtered and processed through to PCR. Negative controls for DNA extractions contained no particulate matter or filters, and PCR negatives contained no DNA template. Analysis of all negative control samples, using the optimized protocols described below, found no evidence of target DNA across all PCR replicates.

#### Water collection, filtration, and DNA extraction

Three liter (L) water samples were collected for all aspects of this study using three sterile, 1 L high-density polyethylene Nalgene^®^ bottles, which were kept on ice in pre-cleaned marine coolers until filtration, which occurred within 24 hours of collection. All water samples were vacuum-filtered using 47 millimeter (mm) 0.8 micron (µm) nylon filters, and used filters were rolled and preserved in 95% ethanol at room temperature. Total eDNA was extracted from filters using the QIAGEN^®^ DNeasy^®^ Blood & Tissue Kit following the Goldberg et al. (2011) protocol incorporating QIAshredder™ spin columns. The qualities of DNA extracts were visualized using 2% agarose gels and the quantities of DNA were assessed using Thermo Fisher Scientific™ NanoDrop™ technology.

#### Droplet Digital PCR assay

Primers were designed to amplify a 100-base pair (bp) fragment of the mitochondrial NADH dehydrogenase subunit 2 (mtDNA ND2) gene in *P. pectinata*, but not in other elasmobranchs that could co-occur with this species in U.S. waters, or in other *Pristis* sawfishes. To design these primers, mtDNA ND2 sequences for *P. pectinata* (GenBank accession no. KP400584.1) and 17 genetically similar exclusion species were downloaded from GenBank (Online Resource 1) and aligned in CodonCode v. 6.0.2 (CodonCode Corporation, Dedham, MA, USA). Forward (PpecND21F: 5’-CTGGTTCACATTGACTCTTAATTTG-3’) and reverse (PpecND21R: 5’-GCTACAGCTTCAGCTCTCCTTC-3’) primers and an internal PrimeTime^®^ double-quenched ZEN™/IOWA Black™ FQ probe labeled with 6-FAM (PpecND2Probe1IBQF: 5’-TACCATAGCCATCATCCCATTATTATTC-3’) were designed to amplify DNA in only *P. pectinata* by including bp differences in the primers and the probe in all exclusion species (see Online Resource 1). To confirm that the combination of the primers and probe amplified the desired locus, PCRs were conducted using quantitative real-time PCR (qRT-PCR) and total genomic DNA (gDNA) from four *P*. *pectinata* individuals. Reaction mixtures contained 1.1 µL of extracted DNA (~25 ng/µL), 1X Bio-Rad^®^ iTaq™ universal probe supermix, 900 nanomolar (nM) of each primer, and 170 nM of probe, adjusted to 22 µL using PCR-grade water. Cycling conditions consisted of enzyme activation at 95°C for 10 min, followed by 40 cycles of: 94°C for 30 s and 64°C for 2 min, followed by enzyme deactivation at 98°C for 10 min, using a ramp rate of 1°C/s. The resulting amplicon from one *P*. *pectinata* individual was cleaned using a QIAGEN^®^ QIAquick PCR Purification Kit using the manufacturer’s protocol, with the exception that all centrifugation steps were conducted at 12,000 rpm for 2 min. Forward and reverse sequences were generated using a BigDye™ Terminator v3.1 Cycle Sequencing Kit (Applied Biosystems™, Foster City, CA, USA) on an Applied Biosystems™ 3730XL DNA Analyzer. A consensus sequence was assembled in CodonCode v. 6.0.2 (CodonCode Corporation, Dedham, MA, USA) and its identity was verified as *P. pectinata* using the NCBI BLAST search function; the generated sequence was 99.3% similar to *P. pectinata* GenBank accession no. KP400584.1 (Chen et al. 2016).

The PCR reaction and cycling conditions were optimized for the Bio-Rad^®^ QX200™ AutoDG™ Droplet Digital™ PCR System (Instrument no. 773BR1456) by systematically adjusting seven variables (i.e., primer and probe concentrations, cycle number, ramp rate, annealing temperature, denaturation time, and elongation time) to produce positive results with high relative florescence units (RFUs) and little to no “droplet rain” (i.e., droplets, or clusters of droplets, that lie between the positive and negative droplet bands on the ddPCR™ scatter plot) (see Online Resource 2). Optimized ddPCR™ reaction mixtures contained 1.1 µL of extracted DNA, 1X Bio-Rad^®^ ddPCR™ supermix for probes (no deoxyuridine triphosphate (dUTP)), 900 nanomolar (nM) of each primer, and 170 nM of probe, adjusted to 22 µL using PCR-grade water. Optimal ddPCR™ cycling conditions were enzyme activation at 95°C for 10 min, followed by 40 cycles of: 94°C for 30 s and 64°C for 2 min, with a final enzyme deactivation step at 98°C for 10 min, using a ramp rate of 1°C/s. To ensure the assay was species-specific for *P. pectinata* in U.S. waters, the optimized ddPCR™ reaction and cycling conditions were tested using 0.20 ng/µL gDNA derived from fin clips from four *P. pectinata* individuals and one individual for each of 12 representative exclusion species (Table 1). The target DNA fragment was amplified in all five ddPCR™ replicates for each *P. pectinata* individual but was not amplified in any of the ddPCR™ replicates for any representative species from five genetically similar ray genera and two shark genera that could co-occur with *P. pectinata*, or in other *Pristis* sawfishes.

**Table 1.**
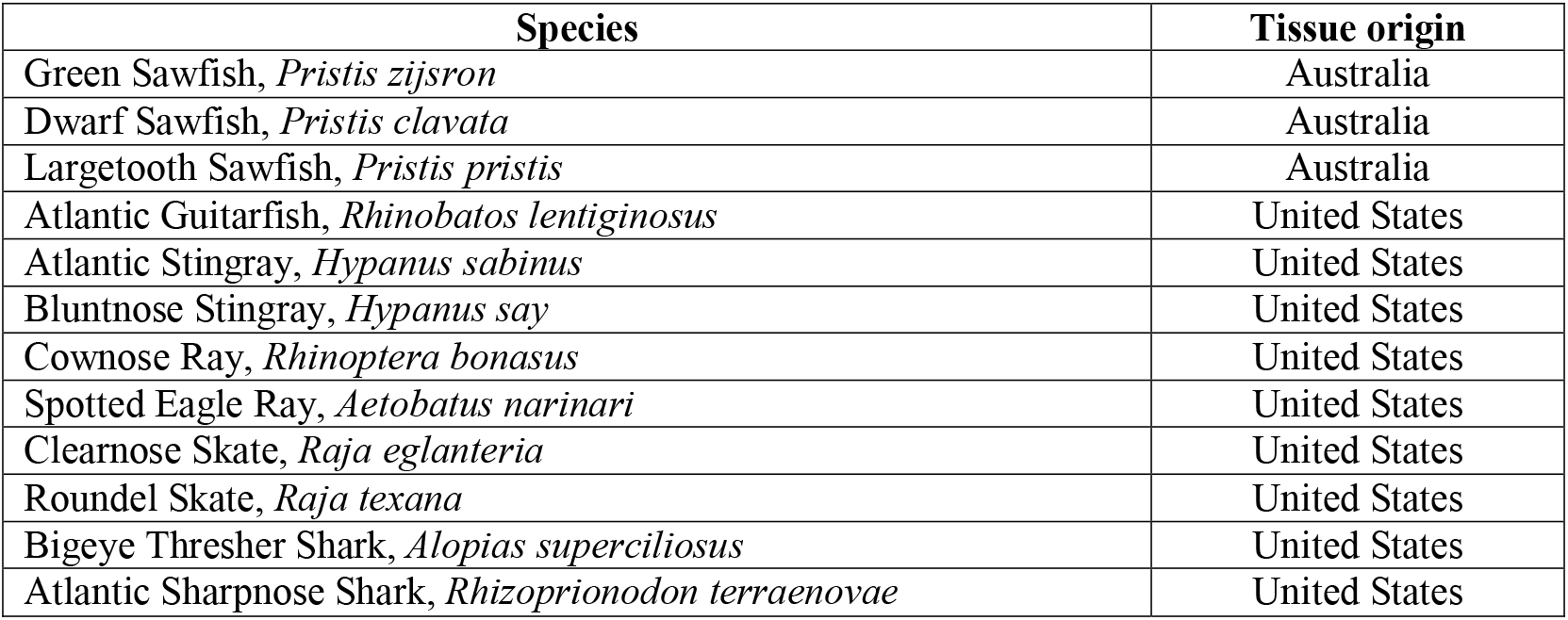
List of 12 exclusion species that were tested to ensure species-specificity of the primers and probe developed for the mitochondrial NADH dehydrogenase subunit 2 (mtDNA ND2) gene in *Pristis pectinata* on the Bio-Rad® QX200™ AutoDG™ Droplet Digital™ PCR System. The country of origin for each tissue sample is included

#### Data analysis

Data were analyzed using three criteria for positive *P. pectinata* detections: 1) droplets fell above a manual threshold (MT) defined for this assay, 2) droplets above the MT fell within the prescribed range of the positive droplet population for this assay (5000–7000 RFUs; Fig. 1), and 3) the concentration of target DNA, determined using Bio-Rad^®^ QuantaSoft™ software using the Rare Event Detection (RED) setting, was at or above the Limit of Detection (LoD) of the assay. Defining an assay-specific MT minimizes the likelihood of incorrectly calling artifact droplets (i.e., droplets that fall above the negative band population in the absence of target DNA; see Online Resource 3) as positive detections (e.g., Hunter et al. 2017). To define an appropriate MT for the *P. pectinata* eDNA assay, a No Template Control (NTC) plate with no target DNA was analyzed on the ddPCR™ platform, using the described reaction and cycling conditions. The highest amplitude of the artifact droplets was 2,700 RFUs; therefore, 3000 RFUs was chosen to adopt a more conservative approach to minimize the risk of a calling a false positive.

**Fig. 1.**
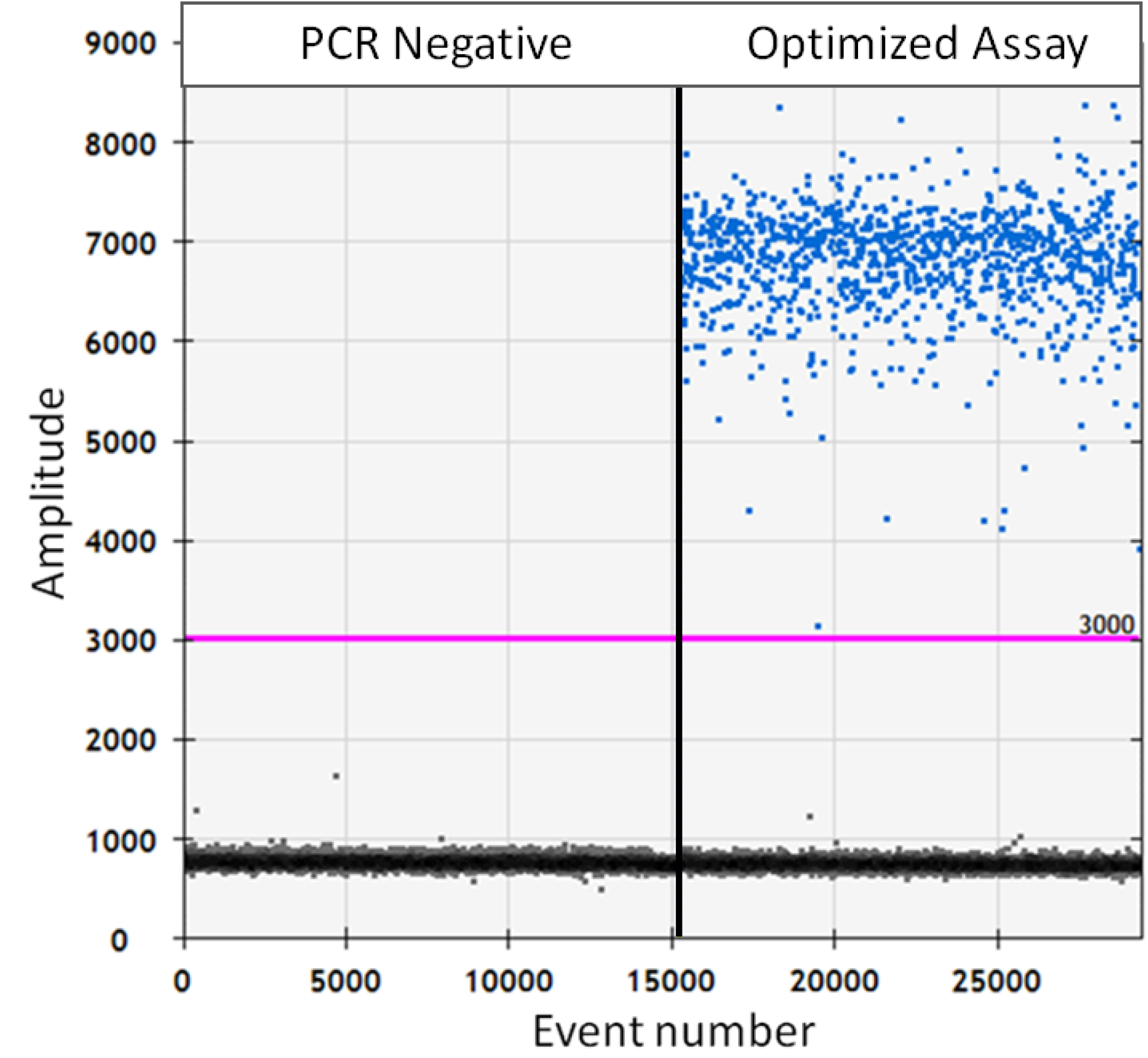
Raw droplet scatter plot of ddPCR™ products using genomic DNA from one Smalltooth Sawfish, *Pristis pectinata*, individual with the optimized assay conditions with a corresponding negative control. Each droplet in each well was classified as either positive (above 3000) or negative (below 3000) for target DNA based on a manual threshold amplitude of 3000 relative florescence units (RFUs); detected using a Bio-Rad^®^ QX200™ Droplet Reader, QuantaSoft™ software and RED analysis setting. Each well is separated by vertical lines, and is labeled to correspond with the sample it represents

To determine the LoD of the assay, ddPCR™ reactions were performed using gDNA from three *P. pectinata* individuals with a 6-fold series of 10X dilutions from starting concentrations of 20 ng/μL (i.e., 1:10 to 1:1,000,000). Target DNA was reliably detected in all replicates for all individuals up to the 1:10,000 dilutions, but not in the 1:100,000 dilutions. The standard error of the 1:1,000,000 also overlapped with zero (Fig. 2a), making detection at this concentration unreliable. To further refine the LoD, ddPCR™ reactions were performed on subsequent 3-fold series of 2X dilutions from the 1:10,000 dilutions (Fig. 2b). The LoD of this assay was found to be the 1:80,000 dilutions, corresponding to 0.25 pg of target DNA in the reaction (Fig. 2c). The standard errors of the 1:80,000 dilutions did not include zero, or overlap with the 1:100,000 dilutions; so using the average number of copies of target DNA/μL in the 1:80,000 dilutions and applying the lower standard error as the relaxed detection threshold (see Baker et al. 2018), the LoD of the assay was determined to be 0.08 copies/μL.

**Fig. 2.**
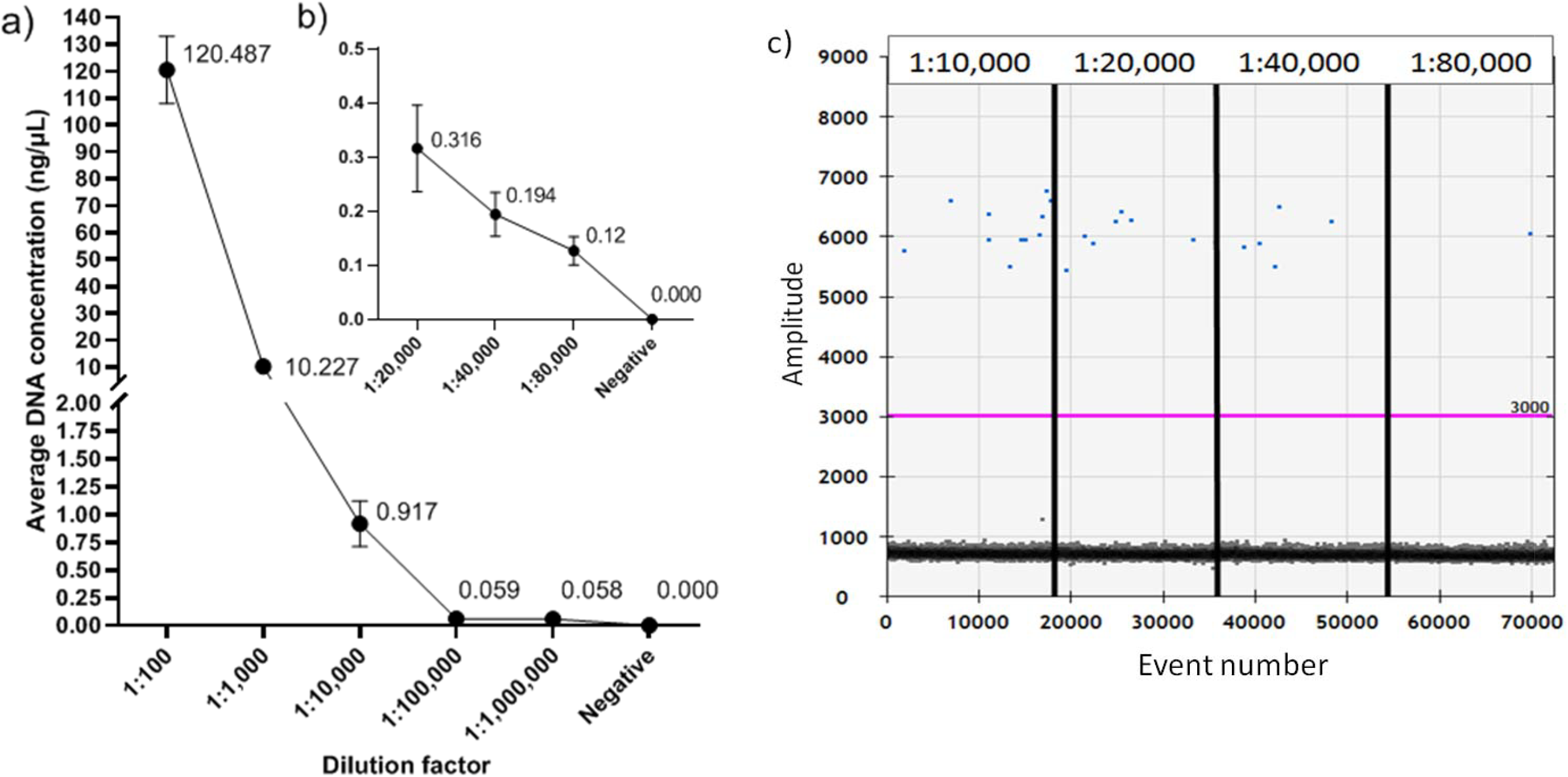
Average target DNA concentrations (copy number/μL) of the Limit of Detection (LoD) dilution series, using genomic DNA from three Smalltooth Sawfish, *Pristis pectinata*, individuals with five replicates each in a) a 6-fold series of 10X dilutions from a starting concentration of 20 ng/µL, b) a 3-fold series of 2X dilutions from the 1:10,000 dilution, and c) a corresponding raw ddPCR™ scatter plot of serial dilution reactions from one replicate of one Smalltooth Sawfish, *Pristis pectinata*, individual. The Bio-Rad^®^ QX200™ Droplet Reader and QuantaSoft™ software using the RED analysis setting was used across all samples, and each droplet in each well was classified as either positive (above 3000) or negative (below 3000) for target DNA based on a manual threshold amplitude of 3000 relative florescence units (RFUs)

### Validation of the *Pristis pectinata* eDNA assay

To validate the ddPCR™ assay, positive *P. pectinata* eDNA samples were acquired via analysis of a water sample from known habitat and through a tank experiment. To collect these positive water samples, a pre-cleaned ~160 L tank was filled with ambient surface water from known *P. pectinata* nursery habitat in the Caloosahatchee River, Florida, approximately 330 m outside of the Harbour Isles Marina. A 3 L water sample was immediately collected from the tank to assess whether *P. pectinata* eDNA was present in the Caloosahatchee River water. One juvenile female *P. pectinata*, measuring 786 mm stretched total length, was captured in a gill net inside the Harbour Isles Marina and placed into the tank. An aerator was added to the tank and dissolved oxygen and water temperature were monitored for the duration of the experiment. A 3 L water sample was collected from the tank immediately after the juvenile was added (time zero) and again after 30 min. All water samples were filtered, DNA extracted, run on ddPCR™, and analyzed using the methods developed in this study.

Applying all three criteria for a positive detection of target DNA, the ddPCR reactions containing DNA extracted from ambient water from the nursery contained an average of 11.54 copies/µL (SE = 0.72) of *P. pectinata* eDNA (Fig. 3). The amount of target eDNA increased to 739.4 copies/µL (SE = 38.31) immediately after the juvenile was added to the tank (time zero) and then increased to 3,175.8 copies/µL (SE = 589.3) after 30 min (Fig. 3). At 30 min, the large quantity of target DNA isolated from the water sample oversaturated the PCR product, resulting in a high standard error.

**Fig. 3.**
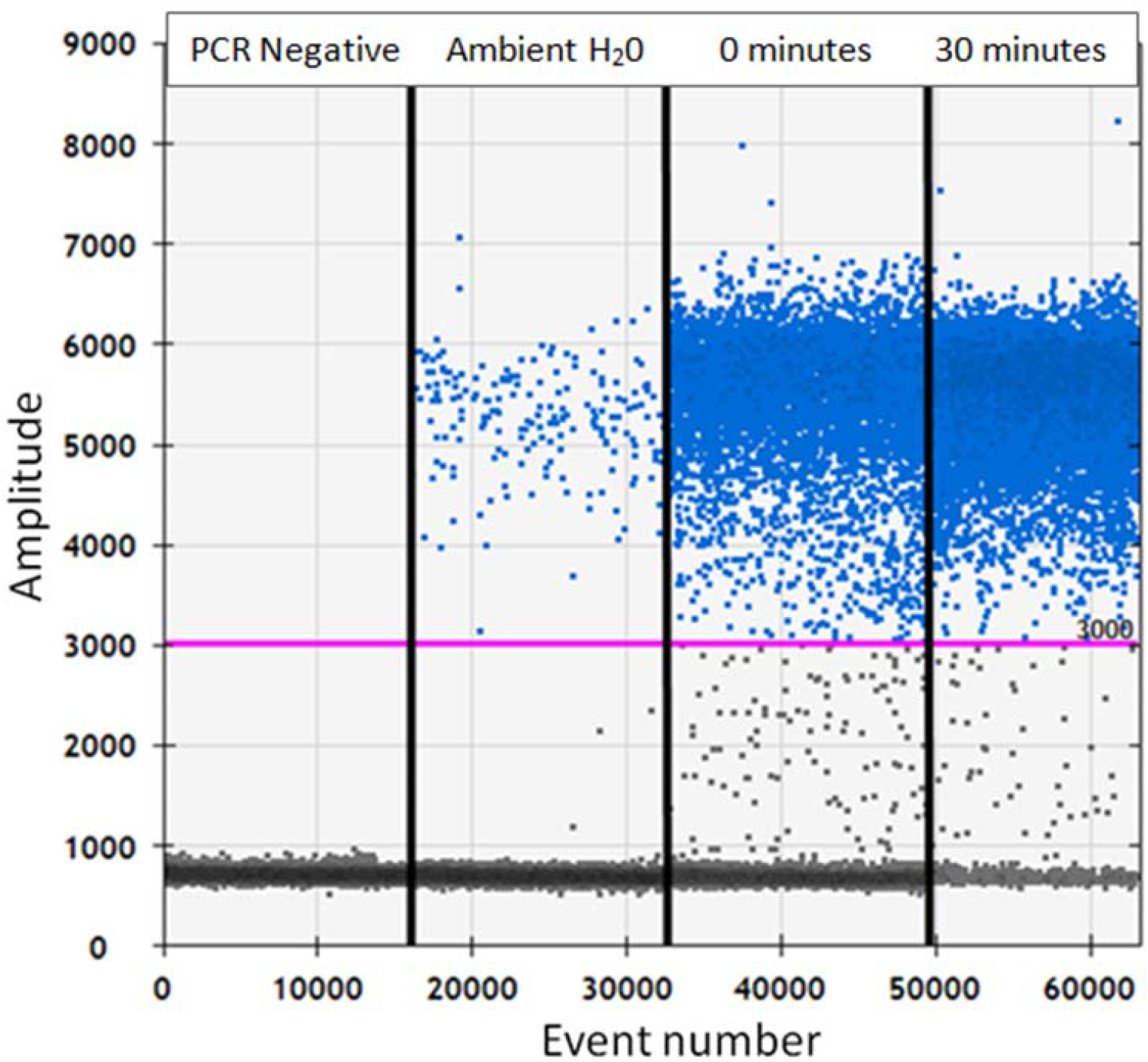
Raw droplet scatter plot of ddPCR™ serial dilution products from one reaction for each of the Smalltooth Sawfish, *Pristis pectinata*, positive eDNA samples. Ambient water refers to water samples collected from the Caloosahatchee River, known *P*. *pectinata* nursery habitat. 0 and 30 min reactions correspond to the positive water samples collected from the *ex situ* tank containing a live *P*. *pectinat*a. Each droplet in each well was classified as either positive (above 3000) or negative (below 3000) for target DNA based on a manual threshold amplitude of 3000 relative florescence units (RFUs) detected using a Bio-Rad^®^ QX200™ Droplet Reader, and QuantaSoft™ software using the RED analysis setting. Each well is separated by vertical lines and is labeled to correspond with the sample or time stage it represents. Note: “Droplet rain” (i.e., droplets, or clusters of droplets, that lie between the positive and negative droplet bands on the ddPCR™ scatter plot) is seen at 30 min due to an oversaturation of target DNA

## Discussion

The developed eDNA assay provides a rapid-assessment tool to conduct targeted surveys to investigate the occurrence and infer the status of *P. pectinata* beyond their contemporary core range. This assay has been validated in the Caloosahatchee River, Florida, where *P. pectinata* is the sole species of sawfish; however, because the assay did not amplify DNA in other *Pristis* sawfishes, it can also be used in locations where other sawfishes have been known to co-occur, at least historically, such as Texas (Brame et al. 2019). It is promising to note that there are no bp differences in the primer or probe sequences designed in this study compared to a mtDNA ND2 sequence for a *P. pectinata* collected in Mexico (GenBank accession no. MF682494.1; Diaz-Jaimes et al. 2018), indicating the developed assay should amplify the target gene in this species from other, nearby waters in the Western Atlantic. Use of the assay outside these waters requires careful consideration and, potentially, further testing. MtDNA genes are often variable among populations within a species (Rubinoff et al. 2006); therefore, before using this assay to conduct eDNA surveys in other geographic regions (e.g., Eastern Atlantic), the primers and probe should be tested with *P. pectinata* tissue samples obtained from the local population. Where fresh *P. pectinata* tissue samples are not available for such testing due to the possibility of local extinctions, historic rostra can be used as an alternative source of DNA (Phillips et al. 2009). Finally, the primers and probe developed here were cross-tested with representative species from closely related genera found in U.S. waters; testing with additional exclusion species is required to ensure that the assay remains species-specific in other geographic regions, highlighting the need for local fisheries knowledge (Poulakis and Grubbs 2019).

The use of ddPCR™ for single species detections is gaining popularity in eDNA research due to its unparalleled ability to detect minute quantities of target DNA amongst high concentrations of non-target DNA and in the presence of natural inhibitors found in water samples (Evans et al. 2017; Hunter et al. 2018). DdPCR™ assays developed for species such as the Bull Shark, *Carcharhinus leucas* (Schweiss et al. In Press) and Killer Whale, *Orcinus orca* (Baker et al. 2018), have found this platform to be capable of detecting less than 0.5 pg of target DNA in a reaction. Such highly sensitive assays are especially critical in eDNA surveys targeting Critically Endangered or Endangered species, where there can be substantial conservation outcomes based on the results of such surveys (Hunter et al. 2018; Poulakis and Grubbs 2019). For instance, the use of ddPCR™ could reduce the risk of false negatives (i.e., where target DNA is present but not detected) stemming from the use of a less sensitive PCR platform such as conventional or qRT-PCR, which are unlikely to detect such minute quantities of target DNA (Doi et al. 2015). Conservation and management strategies developed on the basis of such false negatives could lead to slower implementation and inadequate protections along with incomplete habitat designations for threatened species, ultimately hindering species recovery.

Using the three-criteria approach described here to define positive detections on the ddPCR™ platform provides a rigorous approach to interpret the results of eDNA surveys, reducing the risk of incorrectly calling PCR artifacts as positive species detections (e.g., false positives). For example, using only a MT, an artifact droplet just above the threshold could be incorrectly interpreted as a positive detection. Ensuring the quantity of target DNA is also within the detection capabilities of an assay allows for more robust and confident positive detections. False positives can also result from contamination between eDNA samples or from exogenous DNA. Given the detection capabilities of ddPCR™ assays, strict protocols to prevent contamination (see Goldberg et al. 2016; Schweiss et al. In press) coupled with testing for contamination at every stage in sample processing are critical in producing reliable data from eDNA surveys that may be used as a part of conservation planning. This is especially important when the results of eDNA surveys could be used to prioritize research and management initiatives as well as in the allocation of resources (Poulakis and Grubbs 2019).

With a well-designed water sampling regime, strict field and laboratory controls, and a highly sensitive ddPCR™ assay, targeted species eDNA surveys provide a powerful tool to improve our knowledge of the occurrence of *P*. *pectinata*. The eDNA tool developed here can be used to provide quantitative baseline data in non-core areas from which to measure future progress towards species recovery. Recovery in *P*. *pectinata* populations is predicted to be a slow process due to their life history characteristics and will be dependent on the mitigation of anthropogenic activities (e.g., accidental fisheries mortalities; Carlson and Simpfendorfer 2015). Range re-expansion during recovery is predicted to begin in locations closest to the core population(s) as a result of spillover from adjacent areas, in a stepping-stone fashion (see Saura et al. 2014). There is, however, the possibility that because female *P*. *pectinata* have been shown to exhibit philopatry (Feldheim et al. 2017), occurrence and encounter reports of juvenile *P*. *pectinata* in non-core areas further away from SWFL (e.g. Texas, Mississippi) represent remnant *P*. *pectinata* populations scattered over portions of their former range; under such a scenario, patterns of recovery could be more complex and would ultimately depend on the availability of suitable habitat and the mitigation of threats from anthropogenic activities (Seitz and Poulakis 2006; Poulakis et al. 2011; Norton et al. 2012; Scharer et al. 2017). Conducting targeted eDNA surveys for *P. pectinata* across all historically-occupied regions in U.S. waters could not only aid in conservation planning and prioritizing areas for research, but could also increase our understanding of patterns of recovery in a highly threatened marine species.

## Supporting information

Online Resource 1

Online Resource 2

Online Resource 3

## Acknowledgements

This research was funded by The University of Southern Mississippi and was supported by the Mississippi IDeA Network of Biomedical Research Excellence (INBRE), funded under Institutional Development Award (IDeA) number P20-GM103476 from the National Institutes of General Medical Sciences of the National Institutes of Health. Thank you to Dr. Jonathan Lindner for general advice regarding Droplet Digital™ PCR. Thanks to Joshua Speed, London Williams, and Michael Garrett for laboratory access at University of Mississippi Medical Center (UMMC), and for use of Droplet Digital™ PCR equipment. Thanks to Alia Court and Andrew Wooley for field and laboratory support in Florida. Positive sample collection in Florida was supported by funding from the U.S. Department of Commerce, National Oceanic and Atmospheric Administration’s (NOAA) National Marine Fisheries Service through Section 6 (Cooperation with the States) of the U.S. Endangered Species Act under grant award NA16NMF4720062 to the Florida Fish and Wildlife Conservation Commission. Florida sampling was also supported by Keystone Grant 384 to GRP from the Save Our Seas Foundation. Statements, findings, conclusions, and recommendations are those of the authors and do not necessarily reflect the views or policies of the funders. This research was conducted under Endangered Species Permit number 21043 (GRP) issued by NOAA Fisheries.

