## Supplementary material for "An environmental DNA tool for monitoring the status of the Critically Endangered Smalltooth Sawfish, *Pristis pectinata*, in the Western Atlantic": Online Resource 1

### Electronic Supplementary Material

**Online Resource 1.** List of 17 exclusion elasmobranch species and mitochondrial NADH dehydrogenase subunit 2 (mtDNA ND2) gene GenBank accession numbers that were used to manually design species-specific primers and an internal probe for *Pristis pectinata*. Columns include the number of base pair mismatches between *P. pectinata* and each exclusion species within each primer and probe used in the assay

| **Species** | **GenBank Accession Number** | **Forward primer** | **Reverse primer** | **Probe** |
|---|---|---|---|---|
| Green Sawfish, *Pristis zijsron* | JQ519151.1 | 5 | 3 | 0 |
| Dwarf Sawfish, *Pristis clavata* | KF381507.1 | 4 | 2 | 5 |
| Largetooth Sawfish, *Pristis pristis* | NC_039438.1 | 5 | 3 | 2 |
| Narrow Sawfish, *Anoxypristis cuspidata* | KP233202.1 | 3 | 7 | 3 |
| Common Guitarfish, *Rhinobatos rhinobatos* | JQ518913.1 | 5 | 4 | 5 |
| Southern Stingray, *Hypanus americanus* | JN184288.1 | 7 | 8 | 8 |
| Atlantic Stingray, *Hypanus sabinus* | JQ518787.1 | 6 | 5 | 6 |
| Bluntnose Stingray, *Hypanus say* | JQ518788.1 | 5 | 4 | 5 |
| Roughtail Stingray, *Bathytoshia centroura* | KY90963.1 | 5 | 5 | 6 |
| Pelagic Stingray, *Pteroplatytrygon violacea* | KJ641617.1 | 5 | 5 | 8 |
| Bullnose Eagle Ray, *Myliobatis freminvillii* | JQ518847.1 | 6 | 5 | 5 |
| Cownose Ray, *Rhinoptera bonasus* | JX241056.1 | 8 | 4 | 6 |
| Giant Oceanic Manta Ray, *Mobula birostris* | KM364991.1 | 8 | 7 | 5 |
| Yellow Stingray, *Urobatis jamaicensis* | JQ518941.1 | 7 | 6 | 6 |
| Spotted Eagle Ray, *Aetobatus narinari* | KX151649.1 | 8 | 5 | 4 |
| Clearnose Skate, *Raja eglanteria* | JQ518889.1 | 6 | 5 | 7 |
| Bigeye Thresher Shark, *Alopias superciliosus* | MF374733.1 | 2 | 7 | 4 |
