## Supplementary material for "An environmental DNA tool for monitoring the status of the Critically Endangered Smalltooth Sawfish, *Pristis pectinata*, in the Western Atlantic": Online Resource 2

Title: An environmental DNA tool for monitoring the status of the Critically Endangered Smalltooth Sawfish, *Pristis pectinata,* in the Western Atlantic
Authors: Ryan N. Lehman, Gregg R. Poulakis, Rachel M. Scharer, Katherine E. Schweiss, Jill M. Hendon, Nicole M. Phillips

**Electronic Supplementary Material**

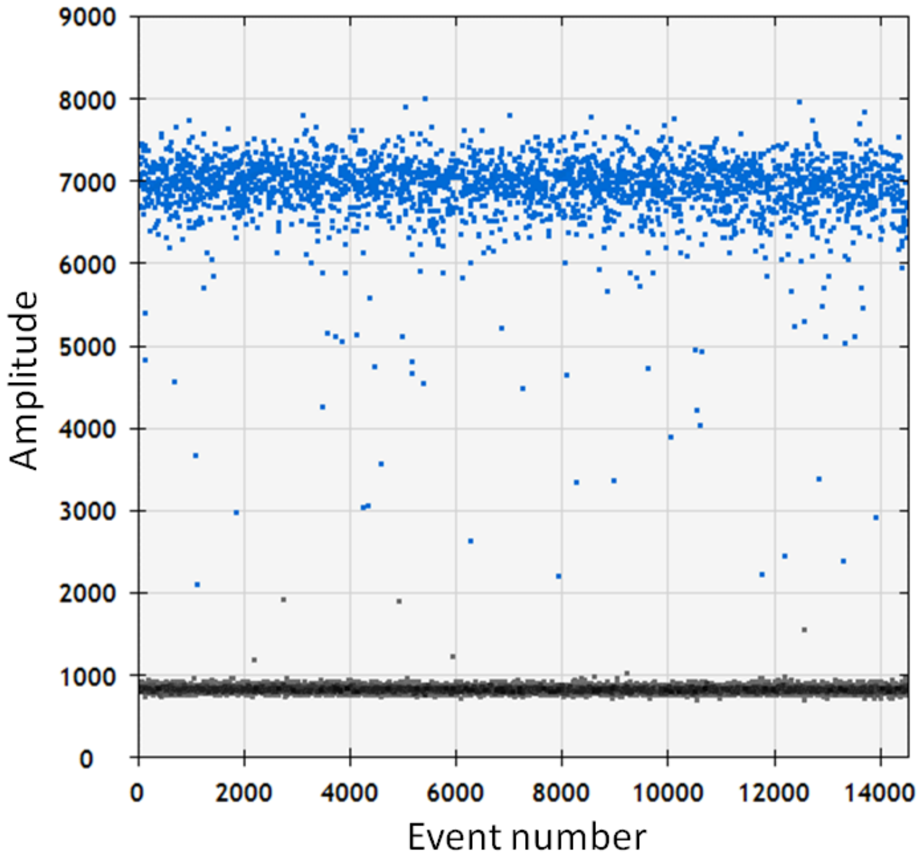


**Online Resource 2.** Raw droplet scatter plot of ddPCR™ products using genomic DNA from one Smalltooth Sawfish, *Pristis pectinata,* individual during assay optimization depicting "droplet rain". Each droplet was classified as either positive (blue) for target DNA, or negative (grey) for target DNA using the Bio-Rad® QX200™ Droplet Reader and QuantaSoft™ software using the RED analysis setting. Because the assay was not fully optimized at this point, "droplet rain" (i.e., droplets, or clusters of droplets, that lie between the positive and negative droplet bands on the ddPCR™ scatter plot) can be seen as the errant droplets that fall outside of the positive droplet population to the manual threshold amplitude of 3000 Relative Fluorescence Units (RFUs)
