## Supplementary material for "An environmental DNA tool for monitoring the status of the Critically Endangered Smalltooth Sawfish, *Pristis pectinata*, in the Western Atlantic": Online Resource 3

**Electronic Supplementary Material**

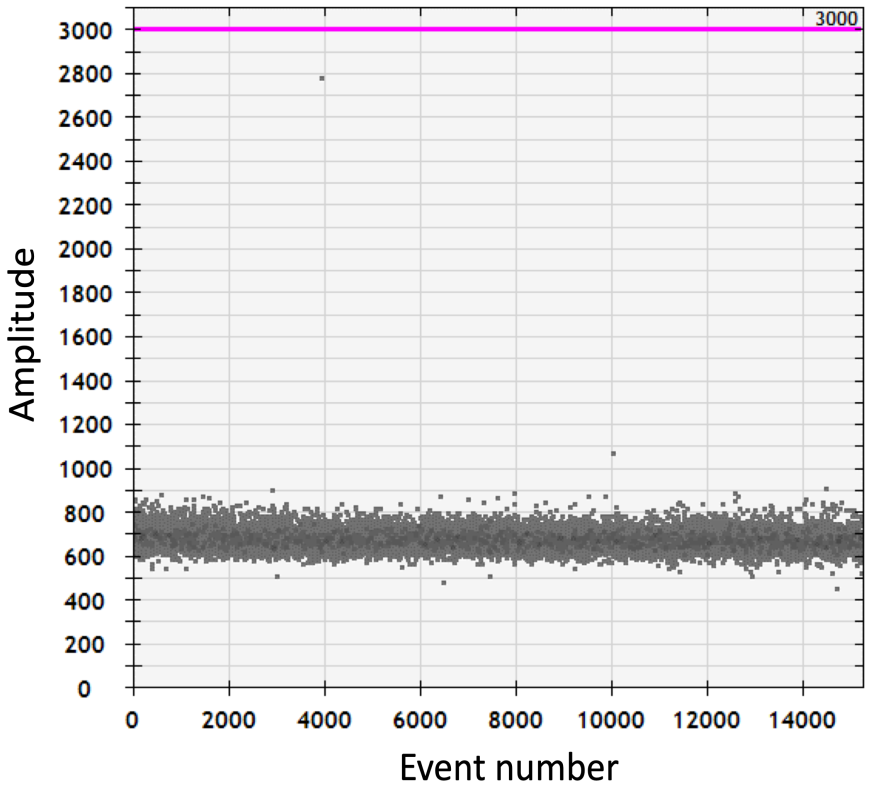


**Online Resource 3.** Raw droplet scatterplot of ddPCR™ products from one replicate of No Template Control (NTC) depicting artifact droplets. Each droplet was classified as negative (grey) due to the absence of target DNA, as detected by the Bio-Rad® QX200™ Droplet Reader and QuantaSoft™ software using the RED analysis setting. Artifact droplets are shown as the errant droplets that fall between the negative droplet population (700–900 Relative Fluorescence Units; RFUs) and the manual threshold amplitude of 3000 RFUs
